# Evaluation of the anti-tumor activities of Sulfonylurea Derivatives

**DOI:** 10.1101/2022.01.11.475798

**Authors:** Sen Lu, Chenchen Guo, Lei Wu, Zhanying Zheng, Xuewen Hua, Wei Wei, Wenqin Zhang, Shaa Zhou, Ruo Li, Sha Zhou

## Abstract

This study prepared 25 sulfonylurea compounds to evaluate anti-tumor activity. Through experimental investigations in MDA-MB-231 and MCF-7, i.e., cell lines of breast carcinoma of human, we have concluded that some compounds can significantly suppress breast carcinoma cells from growing and proliferating. Moreover, the compound M’s inhibitory effect on cells of breast carcinoma is concentration-dependent under a certain treatment time; and the inhibitory effect of the compound M on breast carcinoma cells is time-dependent under a certain concentration. In addition, we also found that the compound M can effectively suppress cells of breast carcinoma from migration and independent survival. The results can show the prospect of research and development of new breast carcinoma treatment drug.

## Introduction

Carcinoma^[1]^ has been found as a major disease that poses a serious threat to human life and health, i.e., malignant tumor^[2]^. Today, breast carcinoma^[3]^ has become a malignant tumor threatening health of females, and the incidences among females are rising year by year. Although numerous treatment methods can be used for breast carcinoma treatment (e.g., surgery^[4]^, radiotherapy^[5]^, chemotherapy^[6]^, targeted drug therapy^[7]^), breast carcinoma, in particular triple-negative breast carcinoma^[8]^, cannot be completely cured because of its strong metastatic ability, and the existing treatment methods have several defects that affect the treatment of breast carcinoma. Thus far, breast carcinoma remains a malignant tumor disease with the highest mortality rate worldwide^[9]^, and it significantly threatens the health of human beings, especially women. Accordingly, new drugs^[10]^ are urgently needed for the treatment of breast carcinoma.

Sulfonylurea compounds have achieved wide applications as hypoglycemic^[11]^ agents in medicine and as pesticides in agriculture^[12]^. Sulfonylurea drugs are found as the earliest but most widely applied oral hypoglycemic drugs, which fall into the first and second generations. As researchers have been investigating the hypoglycemic effect of sulfonylurea drugs, the third generation of sulfonylurea hypoglycemic^[13]^ drugs has been developed. Glimepiride^[14]^ has been found as the main representative of the third-generation sulfonylurea hypoglycemic agents, which shows the advantages of small dosage and the ability to significantly improve insulin resistance. Moreover, Sulfonylurea compounds as herbicides have been investigated as early as the 1970s. After continuous research in the pesticide field, the sulfonylurea herbicides that have been extensively employed currently primarily consist of chlorsulfuron^[15]^, metsulfuron-methyl^[16]^, chlorimuron-methyl^[17]^, etc. As the hypoglycemic mechanism of sulfonylurea compounds has been recently studied in depth, the second-generation hypoglycemic drugs have been found with potential antitumor activity. Thus, the antitumor activity of sulfonylurea compounds has begun to come into the attention in carcinoma treatment. It has become a hot spot in the research of this compound over the past few years.

Pooja Rathore designed and synthesized a series of pyrazoline-substituted benzenesulfonylurea derivatives, and tested 14 compounds in vitro for anti-tumor activity^[18]^. As indicated by the experimental results, these compounds exhibited high malignant tumor inhibitory activity. To be specific, 4 compounds showed broad-spectrum anti-tumor activity and had a strong inhibitory effect on lung carcinoma^[19]^, prostate carcinoma^[20]^, colon carcinoma^[21]^, kidney carcinoma^[22]^, ovarian carcinoma^[23]^ and breast carcinoma^[24]^. A series of 4-phenoxyquinoline derivatives containing sulfonylurea groups were designed, synthesized and assessed for their c-Met kinase inhibitory activity and in vitro cytotoxicity. According to the experimental results, one compound exhibited good selectivity and significant anti-tumor activity against human lung carcinoma cell line HT460, human gastric carcinoma cell line MKN-45, and human colorectal carcinoma cell line HT-29 and the human breast carcinoma cell line MDA-MB-231 with IC_50_ values of 0.055 μM, 0.064 μM, 0.16 μM and 0.49 μM, respectively^[25]^. According to these results, 25 sulfonylurea compounds were prepared to evaluate anti-tumor activity that was reported in the literatures. Moreover, an investigation was conducted on the effect of the new compounds on the migration ability of breast carcinoma cells and the effect on the cloning ability of breast carcinoma cells.

## Material and experiment

### 1. Cell Recovery

The steps of this method took the recovery of a tube of MCF-7 cells as an example.

A tube of MCF-7 cell cryopreservation tube was taken out of liquid nitrogen and quickly put into a 37-degree water bath with float to melt. After the cells completely thawed, they were aspirated with a pipette in the ultra-clean table. Then, they were added to a 15 mL centrifuge tube and pipetted evenly. The 15 mL centrifuge tube was put in the centrifuge at 1000 rpm for 4 min. After the centrifugation, 700 μL of fresh 10% serum DMEM medium was used to blow up the cell pellet, and it was pipetted evenly, and then the cell suspension was averaged. They were divided into 4 wells in a 12-well plate, and then medium was added to each well to make up to 800 μL. Last, the 12-well plate was put in a 37-degree incubator for culture.

### 2. Cell exchange

The steps of this method took the MCF-7 cells in a 12-well plate as an example.

When the cell culture medium in the 12-well plate was found to turn yellow, or the growth and proliferation speed of the cells were found to slow down, the medium should be changed. The steps for cell exchange are as follows. The cell culture solution was aspirated in the 12-well plate, and 800 μL of PBS adherently was added to the wall, the 12-well plate was gently shaken, and the PBS was quickly aspirated. 800 μL of fresh culture medium was added to each well, and the cell culture plate was put into the 37°C cell incubator to continue culturing.

### 3. Cell Passaging

The steps of this method took the passage of MCF-7 cells in a 12-well plate as an example.

When the cells in the 12-well plate grew to about 80% of the area of the well plate, cell passaging operations would be required. The culture medium was aspirated in the 12-well plate, and 800 μL of PBS was added to each well. After the 12-well plate was gently shaken, the PBS was quickly aspirated. 200 μL of 0.25% trypsin was added to each well. When the cells were found to be rounded to large cells falling off, 800 μL of medium was added to each well to stop the digestion, and it was pipetted evenly to form a cell suspension. Next, 700 μL of fresh culture medium was added to each well of the new 12-well plate, and then about 100 μL of cell suspension was added to each well. The 12-well plate was shaken gently and carefully placed in a 37°C cell culture incubator for culture.

### 4. Cell cryopreservation

The steps of this method took a tube of MCF-7 cells cryopreserved as an example.

The old medium was aspirated in the 12-well plate. 800 μL of PBS was added to each well. The well was washed twice, and the PBS was aspirated. 200 μL of 0.25% trypsin was added to each well for digestion. The 12-well plate was placed under a microscope for observation. When many cells fell off, 800 μL of fresh medium was added to stop the digestion and remove the cells, and it was pipetted evenly to form a cell suspension. The 2-well cell suspension was transferred in the 12-well plate to a 15 mL centrifuge tube. Then, the centrifugation was made at 1000 rpm for 4 min. After the centrifugation, the supernatant was aspirated and discarded, and 900 μL of fresh medium was added to blow up the cell pellet and pipette evenly. 100 μL of DMSO was added and mixed quickly. The cell suspension was transferred to the cryopreservation tube and marked. The cryopreservation tube was put into the gradient freezer box, and then the gradient freezer box was placed in the −80°C refrigerator overnight. The next day, the cryopreservation tube was transferred in the gradient cryopreservation box.

### 5. Cell count

Take out the 12-well plate to be counted from the incubator, place it in the ultra-clean table and wash one well of the cells with 800 μL PBS twice. 200 μL of 0.25% trypsin was added for digestion, after terminating the digestion. 10 μL of cell suspension was taken out and placed in a 1.5 mL EP tube, and 90 μL of PBS was added and mixed well. Take a clean cell counting plate, place a cover glass on it, take 10 μL of the cell diluent in the 1.5 mL EP tube and slowly add it from the side of the cover glass to make it uniform distributed under the cover glass. The cell counting plate was placed under the microscope for observation. Density of cell suspension: (number of cells in four large grids/4) × 104 × dilution factor. At this time, the calculated number was the number of cells per milliliter of cell suspension. The total number of cells was the cell density multiplied by the cell volume.

### 6. Compound preparation

An appropriate amount of the new compounds was weighed into a 1.5 mL enzyme-free sterile EP tube, and the name of the compounds was marked. According to the relative molecular mass of the compounds, the volume of DMSO was calculated, and then the DMSO was added to make the final concentration 10 mM. A pipette was used to add the corresponding volume of DMSO and mix evenly, so the compound could be completely dissolved. Put the 1.5 mL EP tube into a 95°C metal bath and heat it for 10 min. Lastly, the compounds solution was stored in the refrigerator-20°C.

### 7. Cell viability assay experiment

. MTT method to determine cell viability experiment

The growth of the cells was observed in the 12-well plate, one of the wells was digested when it grew to an appropriate density. It was transferred to a 15 mL centrifuge tube and diluted to a concentration of 50,000 cells/mL. The 96-well plate was taken out, and 100 μL volume of cell suspension was added to each well of the 96-well plate. Then, it was placed in a 37°C incubator overnight. The medium was changed by aspirating the medium in each well, a new control medium or medium with the corresponding compound concentration was added and placed in a 37°C incubator for cultivation. The medium was changed every day. After the compound was treated for the corresponding time, 15 μL of 5 mg/mL MTT solution was added to each well and incubated for 4 h in a 37-degree cell culture incubator in the dark. The 96-well plate was taken out, and the culture medium was aspirated in each well under dark conditions. Then, 150 μL of DMSO was added to each well to dissolve formazan. At last, it was shaken at a medium speed for 10 min on a shaker. Finally, the absorbance of each well of a 96-well plate was detected at a wavelength of 490 nm.

### 8. Cell Scratch Repair Experiment

In the cell scratch repair experiment, the selected cell line was the human breast cancer cell MDA-MB-231 cell with strong migration ability. The specific experiment operation is presented below.

The new 6-well plate was put into the ultra-clean table. The cells were digested in the 12-well plate and transferred to each well of the 6-well plate evenly. When the MDA-MB-231 cells in the 6-well plate were full, a preparation was made for the scratch operation. A 20 μL pipette was used to insert a yellow pipette tip, perpendicular to the horizontal line drawn with the marker pen, and a vertical line was drawn from top to bottom in the middle of each hole of the 6-well plate to form A cell scratch. The original medium was aspirated in the 6-well plate, and each well was washed with PBS three times to wash away all the cells shed by the cell scratches to facilitate subsequent photographs. The corresponding control medium and serum-free medium with the corresponding concentration of compound were added to each well for processing. A picture of the scratched area was taken under a microscope, and then the 6-well plate was placed in a 37-degree cell incubator for culture. After 24 h, pictures of the scratched area of the cells were taken again under the microscope, and Image J software was used to analyze the width of the scratches before and after the treatment. Then, the migration distance and the migration rate were calculated.

### 9. Cell clone formation experiment

This experimental procedure took the clone formation experiment of MCF-7 cells as an example.

MCF-7 cells Cell counts were performed on the wells treated with different compounds. According to the cell count results, about 1000 cells of each type of treated cells were taken and transferred to a new 6-well plate. The cell culture medium was 2 mL. The 6-well plate was shaken appropriately to make the cells evenly dispersed, and then it was placed in a 37°C cell culture incubator and incubated until the clone was formed. The formed cell clone should contain about 50 cells. 1 mL of 4°C pre-cooled anhydrous methanol was added to each well, and the 6-well plate was put in a 4°C refrigerator and fixed for 15 min. 1 mL of 1% crystal violet solution was added to each well of the 6-well plate and incubated at the ambient temperature for 20 min. After the ambient temperature incubation was completed, the crystal violet solution was aspirated in the well plate and rinsed with ultrapure water several times until no new crystal violet blue liquid flowed out. The 6-well plate was place open in a ventilated place for 30 min. The pictures of the 6-well plate were taken, and then the clones were counted.

## Result and Discussion

### 1. Preliminary screening of anti-breast carcinoma effects of new sulfonylurea compounds

25 compounds with potential anti-tumor activity were assessed by the MTT method in the MCF-7 cell line of breast carcinoma cells for cell viability determination. The compounds above were termed sulfonylurea compounds A-Y.

The specific operation is presented below. The prepared compound mother liquor was taken out and left to melt at the ambient temperature. The compound was dissolved in an appropriate amount of medium to a final concentration of 50 μM, and it was treated in the human breast carcinoma cell MCF-7 cell line. The respective compound was provided with 5 replicate wells. After 72 h, the microplate reader was 490 nm. The absorbance was measured at the wavelength, and the growth inhibition rate of the 25 compounds on the MCF-7 cell line was obtained. Fig. 1 presents the results.

**Fig. 1.**
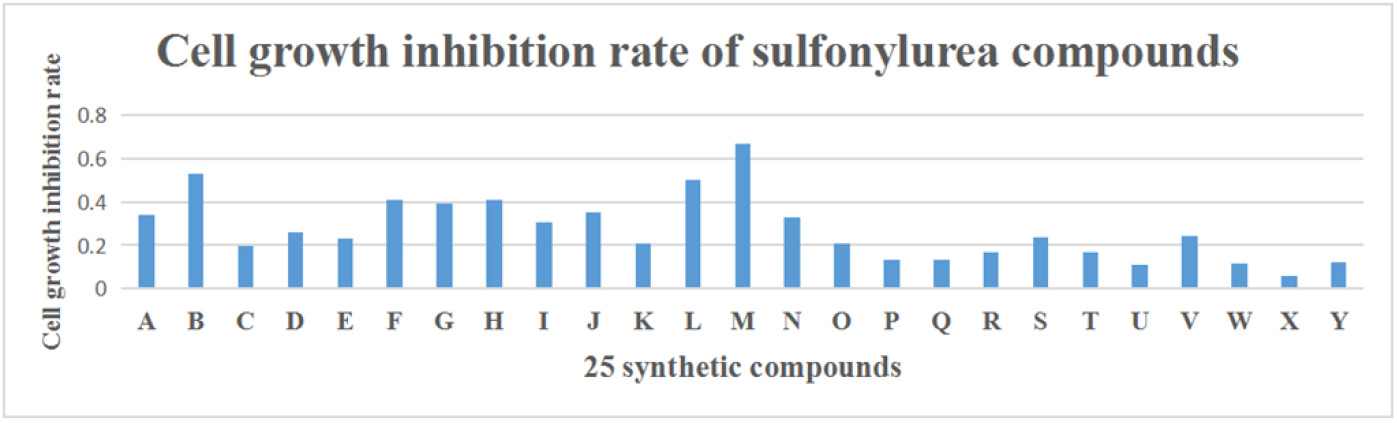
The growth inhibition rate of 25 sulfonylurea compounds at 50 μM on human breast carcinoma cell MCF-7 cell line, the results showed three biological repeats.

As indicated by the experimental results, all compounds exhibited high anti-tumor activity against human breast carcinoma cell MCF-7, especially compounds B, F, G, H, L and M. It is noteworthy that the growth inhibition rate of the sulfonylurea compound M on MCF-7 cells under 50 μM treatment for 72 h was more than 60%, so the antitumor effect of the compound M was demonstrated to be better. Accordingly, in the subsequent research, the compound M would be used as the main research object for several subsequent experiments.

### 2. The inhibitory effect of the concentration gradient of the compound M on breast carcinoma and other tumor cells

#### 2.1 Inhibition of MCF-7 cell line by concentration gradient of the compound M

The compound M with good anti-tumor activity after initial screening was dissolved into the appropriate amount of DMEM cell culture medium. The final concentration was 3.125 μM, 6.25 μM, 12.5 μM, 25 μM, 50 μM and 100 μM compound culture medium. The control was set, and the compound concentration was 0 μM. The MTT method was adopted to determine the concentration of compounds on human breast carcinoma cells MCF-7 cell line the viability of cell survival. The specific operation is presented below. After the cells were planted overnight, different concentrations of compounds were employed to treat the MCF-7 cells every day by changing the medium. After 72 h, the absorbance was measured at a wavelength of 490 nm in the microplate reader, and the cell survival rate was plotted. The line graph with the concentration change and the experimental results are presented in Fig. 2. According to the above experimental results, the inhibitory effect of the sulfonylurea compound M on the human breast carcinoma cell MCF-7 cell line was concentration-dependent.

**Fig. 2.**
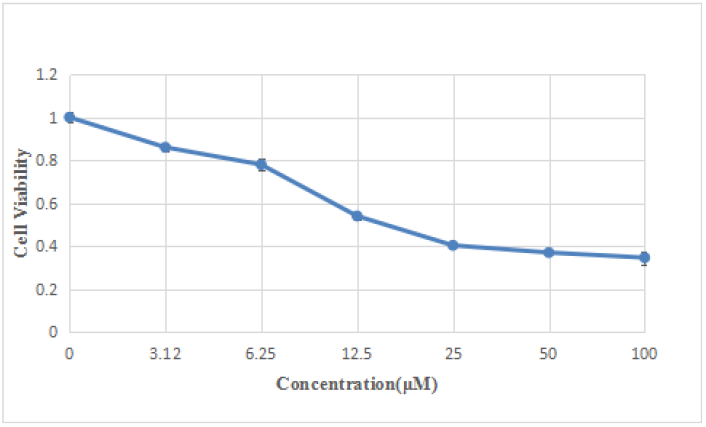
The effect of sulfonylurea M concentration gradient on MCF-7 cell line’s viability, the results showed three biological replicates

#### 2.2 Inhibitory effect of the concentration gradient of the compound M on MDA-MB-231 cell line

The compound M with good antitumor activity was dissolved into the appropriate amount of RPMI-1640 cell culture medium. Their final concentrations were diluted to 3.125 μM, 6.25 μM, 12.5 μM, 25 μM, 50 μM and 100 μM. A control was set. The concentration of the compound was 0 μM. The effect of compounds on cell activity was determined by using MTT method. The specific method is presented below. After MDA-MB-231 cells were planted overnight, 1640 medium with the corresponding compound concentration was used to treat human breast carcinoma cells MDA-MB-231 cells by daily fluid exchange and 72 h later in the enzyme label. The absorbance was measured at 490 nm, and Fig. 3 presents the experimental results. According to the above experimental results, the inhibitory effect of the sulfonylurea compound M on human breast carcinoma cells MDA-MB-231 cells was concentration-dependent.

**Fig. 3.**
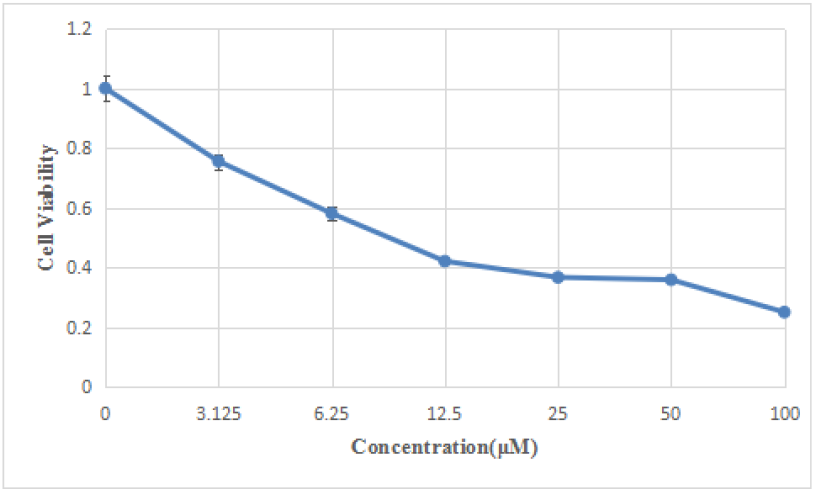
The effect of the sulfonylurea M concentration gradient on the viability of MDA-MB-231 cell line, the results showed three biological replicates.

#### 2.3 Inhibition of SH-SY5Y cell line by the concentration gradient of the compound M

The SH-SY5Y cell line has been found as a human neuroblastoma cell line and a malignant tumor. After the anti-tumor tests on MDA-MB-231 and MCF-7, the cell lines of breast carcinoma of human, it was speculated that the compound M inhibited human neuroblastoma cells. MTT method was applied to verify the conjecture above. The SH-SY5Y cell line was also tested for anti-tumor activity. The concentration gradients of the compound M were still 0 μM, 3.125 μM, 6.25 μM, 12.5 μM, 25 μM, 50 μM and 100 μM and the treatment time was 72 h. Microplate reader detection wavelength was 490 nm. Fig. 4 presents the experimental results of the effects of the compound M on the cell viability of the SH-SY5Y cell line.

**Fig. 4.**
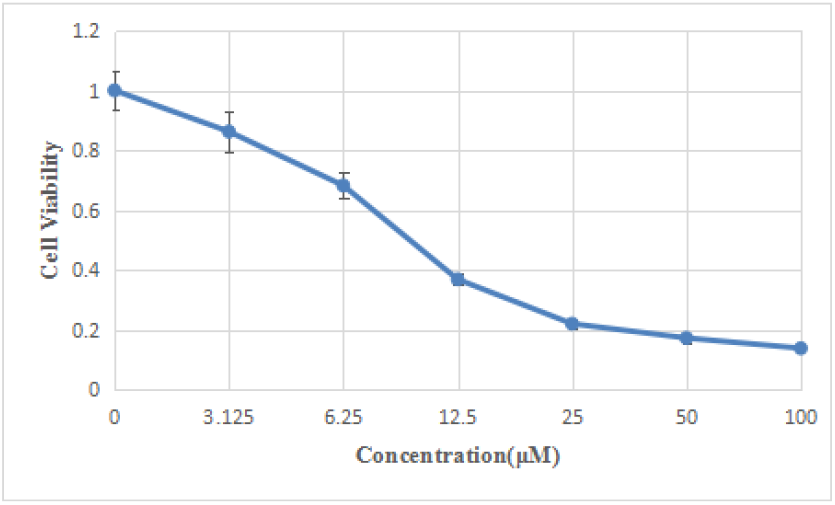
The effect of sulfonylurea M concentration gradient on the viability of SH-SY5Y cell line, the results showed three biological replicates

According to the experimental results, the compound M significantly inhibited the growth of human neuroblastoma cell lines, and as the concentration of the compound increased, the growth inhibitory effect was improved, and the cell viability decreased. Thus, the inhibitory effect of the sulfonylurea compound M on SH-SY5Y cell line was concentration-dependent. Moreover, the experiment proved that the compound M inhibited the growth of human breast carcinoma cells and had a potential anti-tumor effect on other tumor cells, which should be further studied experimentally in depth.

#### 2.4 Calculation of IC_50_ of the compound M in three tumor cells

The half-inhibitory concentration is the concentration of the compound when the inhibitory effect of the compound in tumor cells is 50%. In addition, the half-inhibitory concentration is of high reference significance for investigating the antitumor activity of the compound. Graphpad Prism software was adopted to perform certain statistics and calculations on the half-inhibitory concentration (IC_50_) of the inhibitory effect of the compound M on the three tumor cells. Table 1 lists the relevant results.

**Table 1.** IC_50_ of the compound M in tumor cells

| Compound M | $IC_{50}$ (μM) | | |
| --- | --- | --- | --- |
|  | MCF-7 | MDA-MB-231 | SH-SY5Y |
|  | 8.15±0.46 | 12.25±1.05 | 10.32±0.38 |

**Table 2.**
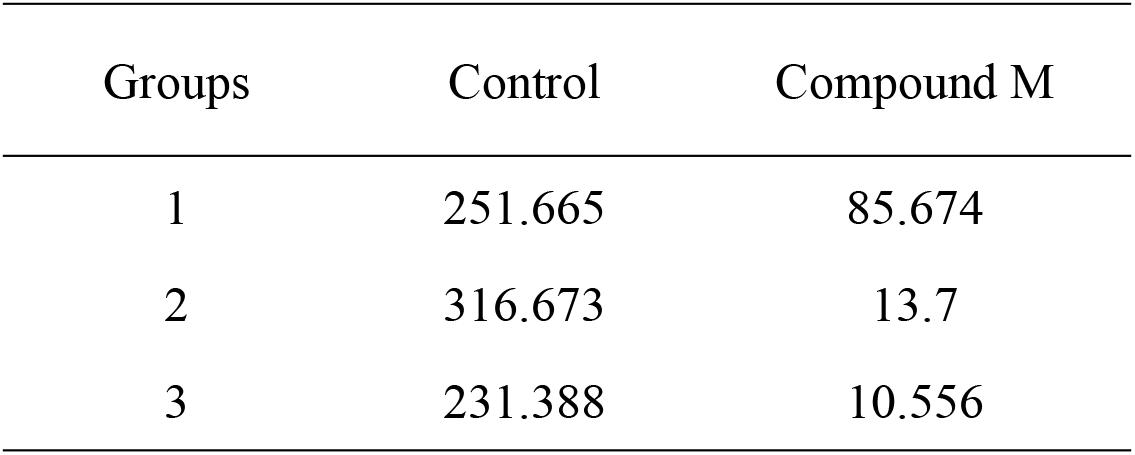
The effect of the compound M on the migration distance (μM) of MDA-MB-231 cell line treated with 100 μM for 24 h (the results showed three biological replicates)

| Groups | Control | Compound M |
| --- | --- | --- |
| 1 | 251.665 | 85.674 |
| 2 | 316.673 | 13.7 |
| 3 | 231.388 | 10.556 |

**Table 3.** The effect of the compound M on the migration rate of MDA-MB-231 cell line treated with 100 μM for 24 h (the results showed three biological replicates)

| Groups | Control | Compound M |
| --- | --- | --- |
| 1 | 0.284 | 0.0885 |
| 2 | 0.36 | 0.015 |
| 3 | 0.288 | 0.0123 |

**Table 4.** Clonal formation experiments of 100 μM compound M treated human breast carcinoma cell line MCF-7

| Groups | Control | Compound M |
| --- | --- | --- |
| 1 | 375 | 231 |
| 2 | 302 | 187 |
| 3 | 333 | 192 |

**Table 5.** Clonal formation experiments after 100 μM compound M treated human breast carcinoma

| cell line MDA-MB-231 |  |  |
| --- | --- | --- |
| Groups | Control | Compound M |
| 1 | 193 | 93 |
| 2 | 201 | 61 |
| 3 | 223 | 80 |

According to Table 1, IC_50_ of the compound M in tumor cells was relatively small, basically around 30 μM and below. This indicated that the compound M could have high tumor growth inhibitory activity, and we are interested in conducting in-depth anti-tumor effects research.

### 3. The inhibitory effect of the compound M on breast carcinoma cells with time gradient

#### 3.1 Inhibition of the compound M time gradient on MCF-7 cell line

The previous experiments reported that the inhibitory effect of the compound M on breast carcinoma cells was concentration-dependent. In other words, in a certain concentration gradient, the compound’s inhibitory effect on breast carcinoma cells increased with the increase in the concentration, and the cell viability decreased with the increase in the concentration.

To further evaluate the anti-tumor effect of the compound M, we fixed the concentration of the compound to 100 μM with better anti-tumor effect, and then set 5 time points in the compound treatment time, respectively 0 h and 24 h, 48 h, 72 h and 96 h, the anti-tumor effects of the compound M were tested using MTT method at different time points to investigate the effect of the time gradient of the compound treatment on the viability of tumor cells. The use of human breast carcinoma MCF-7 cell line was first studied. The specific operation is presented below. After breast carcinoma cells were planted overnight, they were treated with the compound M concentration of 100 μM, with the time points of 0h, 24 h, 48 h, At 72 h and 96 h. The absorbance was measured using a microplate reader. The experimental results are presented in Fig. 5.

**Fig. 5.**
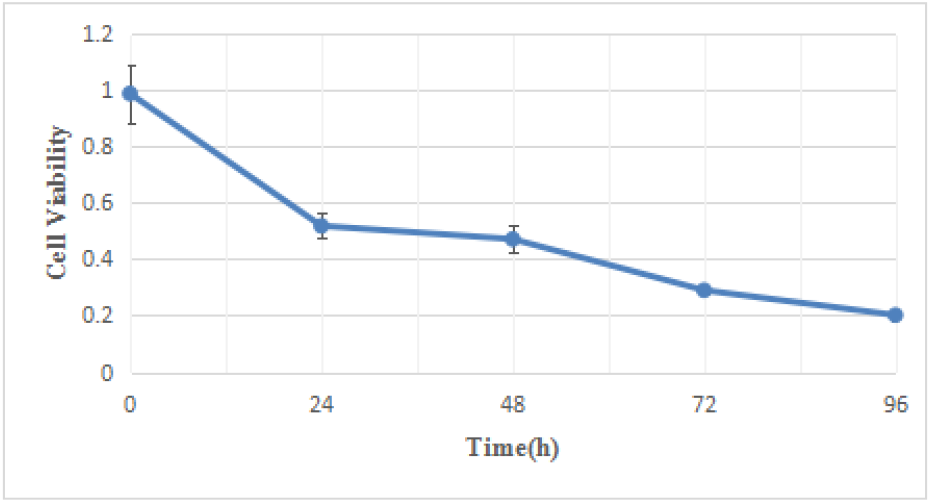
Effect of time gradient of the sulfonylurea compound M on MCF-7 cell line’s viability, the results showed three biological replicates

As indicated by the above results, the inhibitory effect of the compound M on MCF-7 cell line was time-dependent. Under a certain compound concentration, as the treatment time of the compound was extended, the inhibitory effect of the compound on the MCF-7 cell line tended to increase, and the cell viability tended to decrease. After the treatment time was 4 days, the survival rate of breast carcinoma MCF-7 cell line was below 20%.

#### 3.2 Inhibitory effect of the compound M time gradients on MDA-MB-231 cell line

Using the same method as 3.1, the concentration of the compound was still fixed at 100 μM, and the time gradient was still 0 h, 24 h, 48 h, 72 h, and 96 h. Next, the effect of the viability of the MDA-MB-231 cell line was found. Fig. 6 presents the experimental results.

**Fig. 6.**
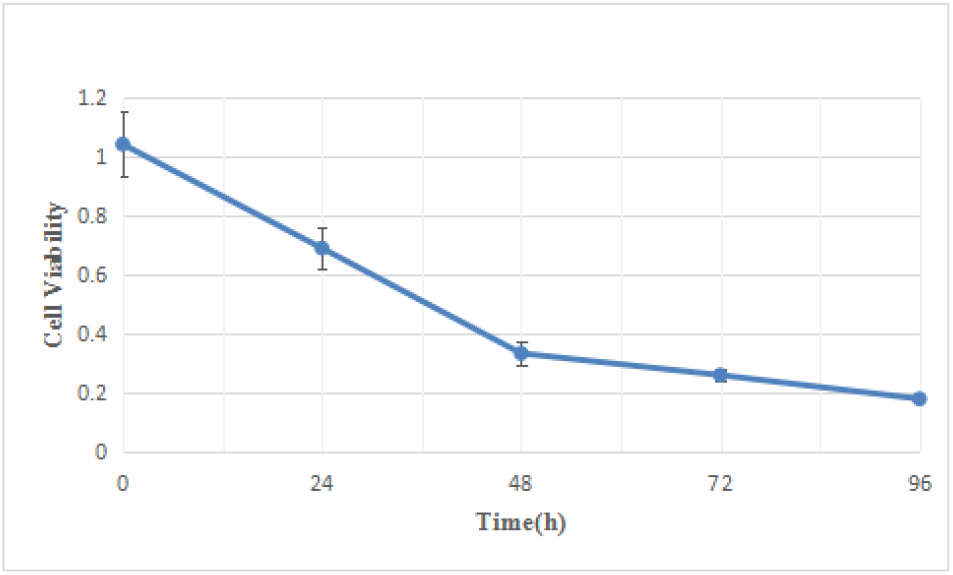
The effect of sulfonylurea M time gradient on the viability of MDA-MB-231 cell line, the results showed three biological replicates

The above experimental result indicates that the growth inhibitory effect of the compound M on the human breast carcinoma MDA-MB-231 cell line is also time-dependent. Under a certain concentration of the compound, as the treatment time increases, its inhibitory effect on tumor cells is enhanced, and the cell viability is reduced.

### 4. The effect of the compound M on the migration ability of breast carcinoma cells

A wound healing assay was designed to investigate the effect of the compound M on the migration ability of tumor cells. During the wound healing assay, the human breast carcinoma MDA-MB-231 cell line with a significant cell migration ability was selected. The specific operation is presented below. The compound M was diluted with the culture medium to make the final concentration of 100 μM. It is noteworthy that a serum-free medium was used to avoid the effects of cell proliferation. Next, the cells were cultured in a 6-well plate. A scratch test was performed after the cells were overgrown. Immediately after scratching, a picture was taken under the microscope, during which the cell scratch width was 0 h. After that, the prepared 100 μM culture medium was added for treatment and placed in a cell incubator for cultivation. After 24 h, the photograph was taken again. Then, the width of the cell scratch after 24h of the compound treatment was taken. In this experiment, a control was set, the compound-free serum-free medium was added, and the other group was the compound M-added experimental group. Fig. 7~10 present the scratch test results of the control and the experimental group. The cell scratch experiment was repeated three times to count and analyze the cell migration distance.

**Fig. 7.**
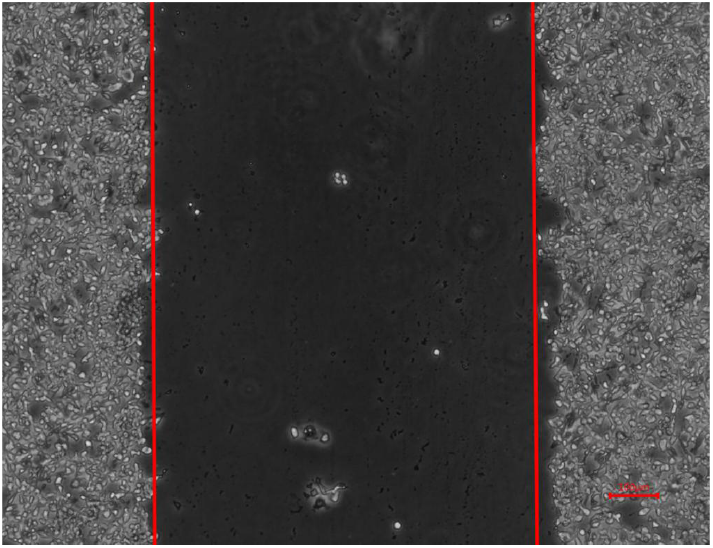
Schematic diagram of cell scratches in the control at 0 h. The distance between the red lines could be considered the scratch width. The scale bar in the figure represents 100 μm.

**Fig. 8.**
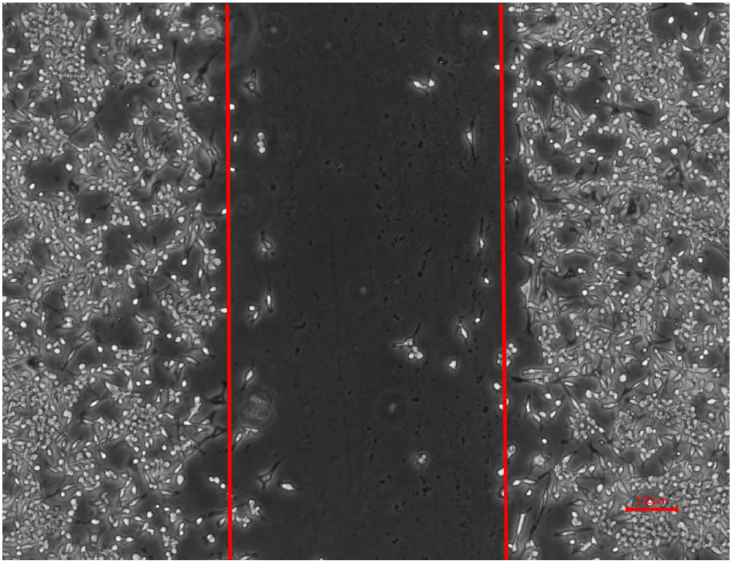
Schematic diagram of cell scratches in the control at 24 h. The distance between the red lines could be considered the scratch width, and the scale bar in the figure represents 100 μm.

**Fig. 9.**
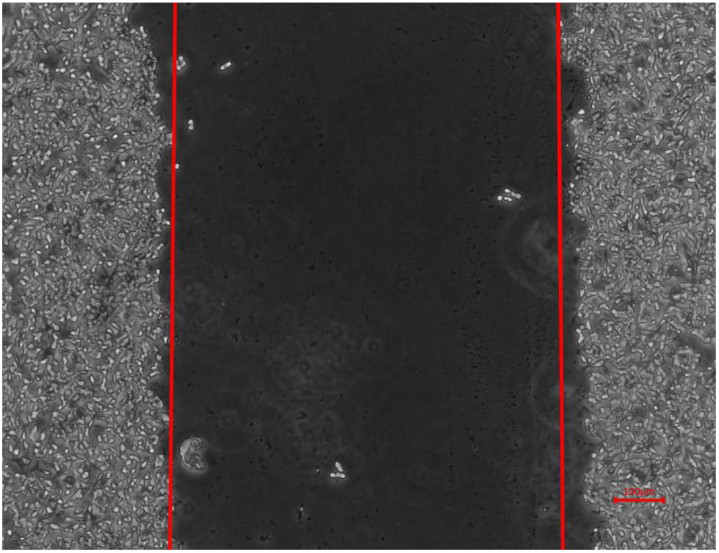
Schematic diagram of cell scratches treated with the sulfonylurea compound M for 0 h. The distance between the red lines could be considered the scratch width, and the scale bar in the figure represents 100 μm

**Fig. 10.**
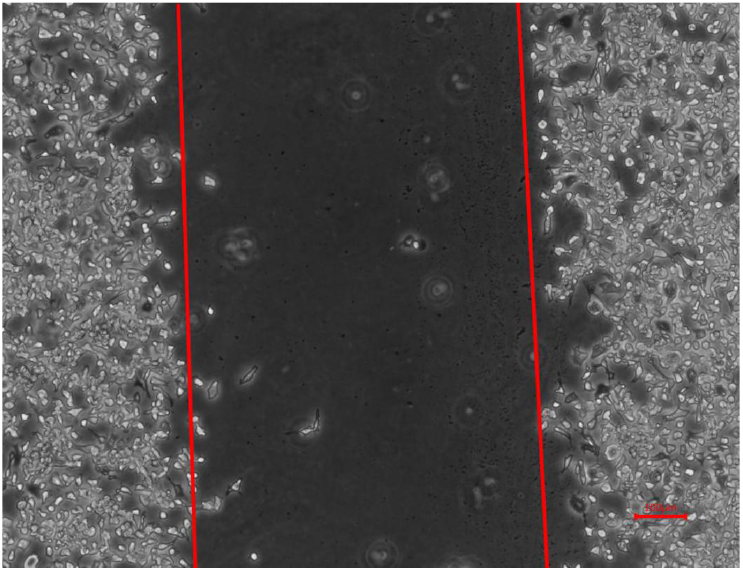
Schematic diagram of cell scratches treated with the sulfonylurea compound M for 24 h. The distance between the red lines could be considered the scratch width. The scale bar in the figure represents 100 μm

**Fig. 11.**
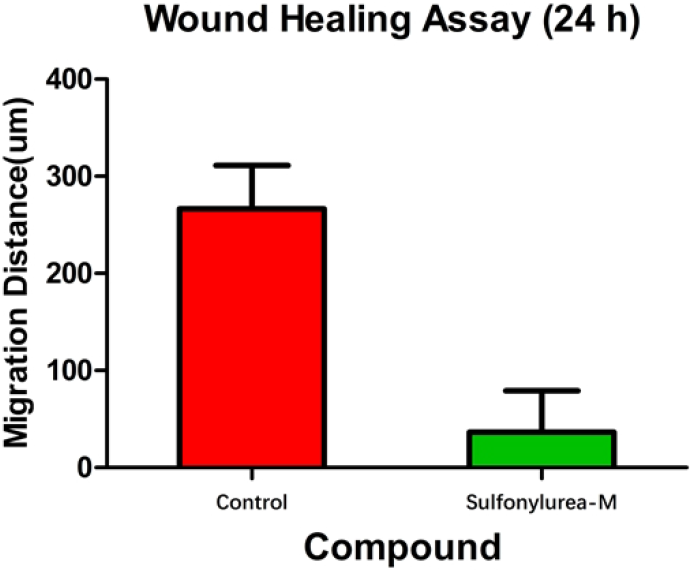
The effect of three new compounds on the migration distance of MDA-MB-231 cell line, the results showed three biological repeats.

As indicated by the experimental results of the wound healing assay above, after 24 h of culture in the serum-free medium of the control, the cells on both sides migrated significantly to the scratch area. Besides, in the experimental group, after 100 μM of the sulfonylurea compound M was used for the treatment, the migration of human breast carcinoma cells MDA-MB-231 was significantly inhibited. To more accurately and intuitively show the effect of the compound M on the migration ability of human breast carcinoma cell MDA-MB-231 cell line, the software Image J was used to count the migration distance of cells in accordance with the scale in the figures (migration distance = 0 h cells Scratch width-24 h cell scratch width), and the mobility was determined (mobility = migration distance / 0 h cell scratch width). The results showed three biological replicates.

As indicated by the above results, the sulfonylurea compound M could significantly reduce the migration ability of human breast carcinoma cell MDA-MB-231 cell line. Breast carcinoma cells MDA-MB-231 had the strongest migration inhibition ability. As shown in Fig. 12, the migration rate of cells in the control was over 30% after treatment with serum-free medium for 24 h, and after 24 h of compound treatment, the migration rate of human breast carcinoma cells could be reduced to about 10%. Thus, through wound healing assay, we can conclude that the compound M not only has a good inhibitory effect on the growth and proliferation of human breast carcinoma cells, but also possesses a strong suppression effect on migration ability of MDA-MB-231 cell line. This shows that the compound M has certain development prospect in the treatment of human breast carcinoma metastasis, but further research and evaluation are needed.

**Fig. 12.**
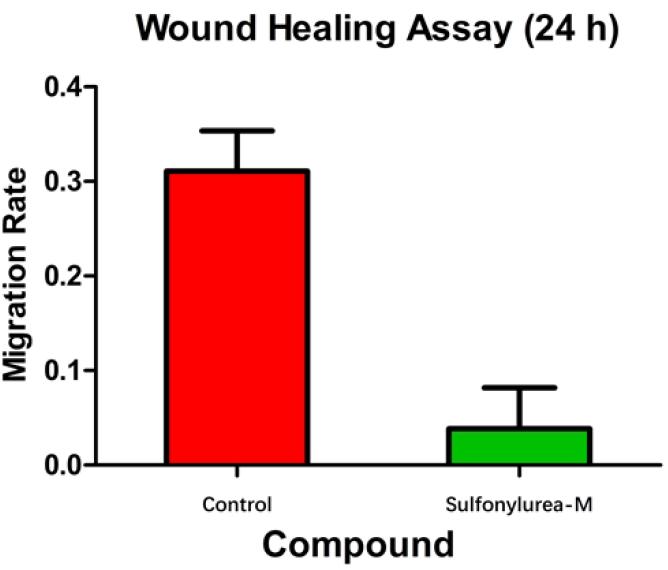
The effect of three new compounds on the mobility of MDA-MB-231 cell line, the results showed three biological replicates

### 5. The effect of the compound M on the cloning ability of breast carcinoma cells

#### 5.1 Inhibition of the compound M on the cloning ability of MCF-7 cell line

Cell clone formation experiments were designed to further study the effect of the compound M on the proliferation and independent viability of tumor cells. Clone formation is one of the effective methods to measure the cell proliferation ability. A single cell was propagated for over 6 generations in vitro, and the cell group composed of its progeny was termed clone. At this time, the number of cell clones was more than 50, and the size was nearly 0.5mm. Based on the counted clones and the calculated clone formation rate, a quantitative analysis of the proliferation potential of a single cell was conducted to understand the proliferation ability and independent viability of the cells.

In the experiment exploring the effect of the compound M on the formation of MCF-7 clones in human breast carcinoma cells, the concentration of the compound we selected was 100 μM with high antitumor effect. The experimental group had three wells, i.e., three biological replicates. The specific operation is presented below. 1000 cells without any compound treatment were added to three wells of 6-well plate. 1000 cells with the treatment of 100 μM sulfonylurea compound M for 24 h were added to three wells, and another 6-well plate was taken. The cells were placed in a cell incubator and incubated. They were observed once every three days until the clones were formed.

After the clones were formed, the cells were fixed, stained and washed with water. In addition, the 6-well plate was placed in a ventilated place for 30 min before taking pictures and counting the clones. Fig. 13 presents the experimental results.

**Fig. 13.**
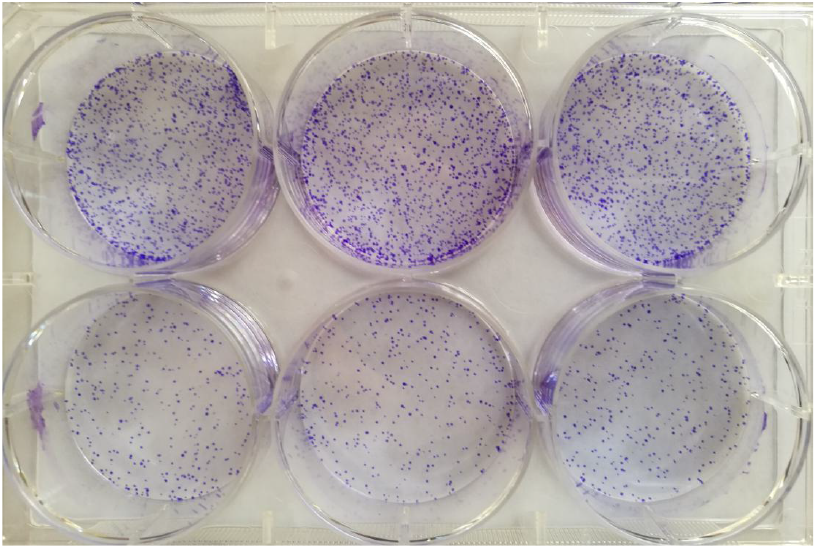
The control and the sulfonylurea compound M treatment clone formation experiment, the upper three holes are the control, the lower three holes are the sulfonylurea compound M treatment group

**Fig. 14.**
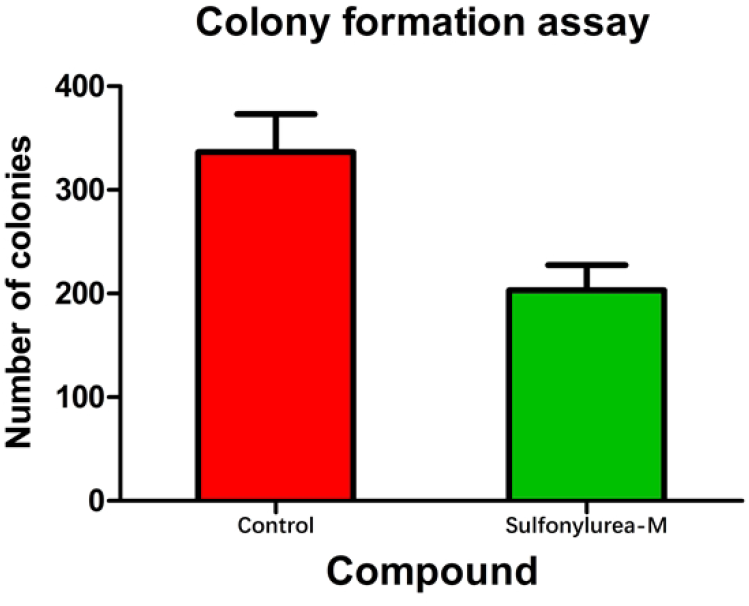
The effect of three compounds on the cloning ability of MCF-7 cells, the results showed three biological replicates

**Fig. 15.**
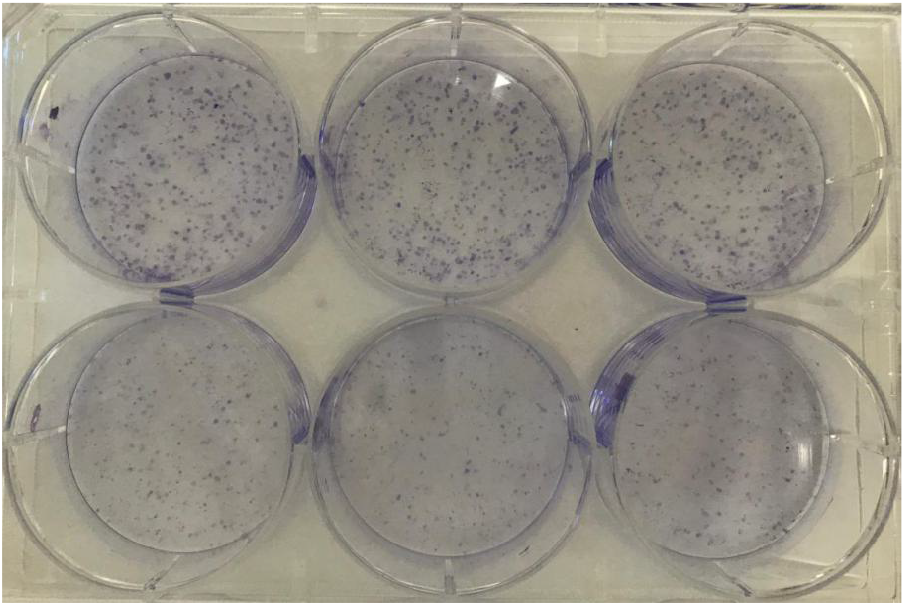
The control and the sulfonylurea compound M treatment after cloning formation experiment, the upper three holes are the control, the lower three holes are the sulfonylurea compound M treatment group

**Fig. 16.**
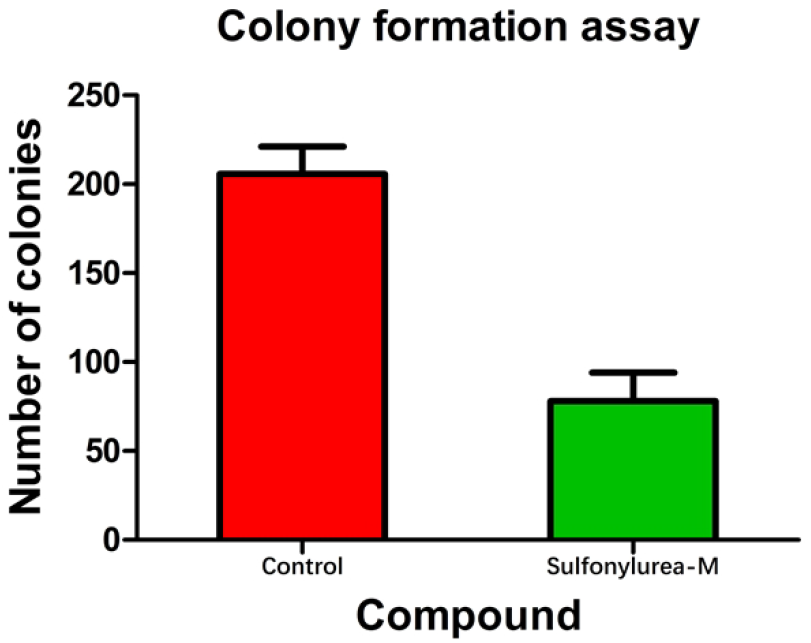
The effect of the compound M on the cloning ability of MDA-MB-231 cells, the results showed three biological replicates

As indicated by the above experimental results, after treatment with the compound M, the human breast carcinoma cell line MCF-7 cells significantly reduced the number of cell clones compared with the control. Accordingly, it was initially concluded that the compound M can significantly inhibit the cloning ability of human breast carcinoma cells MCF-7, greatly reducing the proliferation and viability of MCF-7 cells. For this reason, it can be speculated that the compound M may be able to inhibit breast carcinoma cells from forming malignant tumors for human body. 5.2 Inhibition of the compound M on the cloning ability of MDA-MB-231 cell line

The above results show that the compound M has a good inhibitory effect on the cloning ability of the human breast carcinoma cell line MDA-MB-231, which is basically consistent with the experimental result of MCF-7 cell.

## Conclusion

In this work, we explored the evaluation of anti-tumor activities of 25 sulfonylurea structures. The experiment results showed that compound M can suppress breast carcinoma cells from growing and proliferating. Compound M’s inhibitory effect on cells of breast carcinoma is concentration-dependent under a certain treatment time as well as time-dependent under a certain concentration. It was worthy noting that M can effectively suppress cells of breast carcinoma from migration and independent survival. The study will provide a valuable clue for innovation of new breast carcinoma treatment drug.

## Acknowledgments

The authors would like to thank Xinyi Lu for advice on cell experiments.

